# Network Pharmacology Integrated Molecular Docking Based Prediction of Active Compounds and Potential Targets in *Tinospora crispa* Linn. as Insulin Sensitizer

**DOI:** 10.1101/2021.05.05.442728

**Authors:** Ummu Mastna Zuhri, Erni Hernawati Purwaningsih, Fadilah, Nancy Dewi Yuliana

## Abstract

Insulin resistance is a metabolic disorder characterized by the decreased response to insulin in muscle, liver, and adipose cells. The normal insulin levels are unable to control glucose, lipids, and energy homeostasis. This condition remains a complex phenomenon that involves several genetic defects and environmental stresses, such as obesity. A full understanding is required to understand the entire itinerary and functional consequences of the occurrence of insulin resistance to develop a potent drug in diabetes management. In the present study, we investigated the mechanism of known phytochemical constituents of *Tinospora crispa* and its interaction with insulin resistant target proteins by using network pharmacology and molecular docking. The insulin sensitizer activities of *Tinospora crispa* may be associated with the inhibition of the activation of the inflammatory pathway and the activation of insulin signaling. Tinoscorside A, Makisterone C, Borapetoside A and B, and β sitosterol consider the main phytoconstituents of *Tinospora crispa* by its binding with active sites of main protein targets of insulin resistance potential therapy. In conclusion, *Tinospora crispa* was one of the promising therapeutic agent in type 2 diabetes mellitus management. Regulation in glucose homeostasis, adipolysis, cell proliferation, and antiapoptosis may be the critical mechanism of *Tinospora crispa* as an insulin sensitizer.

## Introduction

Insulin resistance is a metabolic disorder characterized by the decreased response to insulin in muscle, liver, and adipose cells. The normal insulin levels are unable to control glucose, lipids, and energy homeostasis. This condition remains a complex phenomenon that involves several genetic defects and environmental stresses, such as obesity. A full understanding is required to understand the entire itinerary and functional consequences of the occurrence of insulin resistance to develop a potent drug in diabetes management^1^. Furthermore, drug monotherapy is considered difficult to provide the desired effect in controlling blood glucose levels in type 2 diabetes patients. Combination therapy which has different mechanisms is needed for maintaining the requirement of therapeutic management. However, this strategy meets some disadvantages regarding the increasing drug side effects, toxicity, and interactions between the drugs. Another alternative is the search for a drug molecule that selectively modulates different targets and improving the balance of efficacy and safety compared to single target agents^2^.

This approach is in line with the paradigm that has begun to change in natural product drug discovery, the “herbal shotgun” which utilizes the synergy of multi constituents which have varied targets^3^. Until now, research on antidiabetic drugs from *Tinospora crispa* (*T. crispa*) are still using the “silver bullet” approach which is oriented towards the single active principle with one target. As the result, the information obtained was not comprehensively explaining the activity of *T. crispa*, yet it closer to the activity of certain metabolites contained in *T. crispa*^4–12^. Research on medicinal plants conducted by Wink *et al* also supports the herbal shotgun approach. It was reported that the combination of metabolite compounds in the medicinal plant had a better effect than the single isolate of metabolite compound^13^. This fact requires more recent research using the herbal shotgun approach so that the activity of medicinal plants can be comprehensively reveal^3^.

*T. crispa* as a multi-constituent preparation has not clearly known its action mechanism in insulin sensitization activity. It needs to be explored more extensively using a comprehensive approach such as *in silico* prediction using network pharmacology analysis. The current development of bioinformatics technology allows researchers to predict the action mechanism of medicinal plant constituents as multicomponent simultaneously as a therapeutic activity.

In our study, the potential effect and mechanism of *T. crispa* as multi-compounds preparation were analyzed using network pharmacology to find out its activity as an insulin sensitizer. The most significant protein targets then verified its binding with T. crispa constituents by docking molecular strategy to verify its interactions (Fig 1).

**Fig 1.**
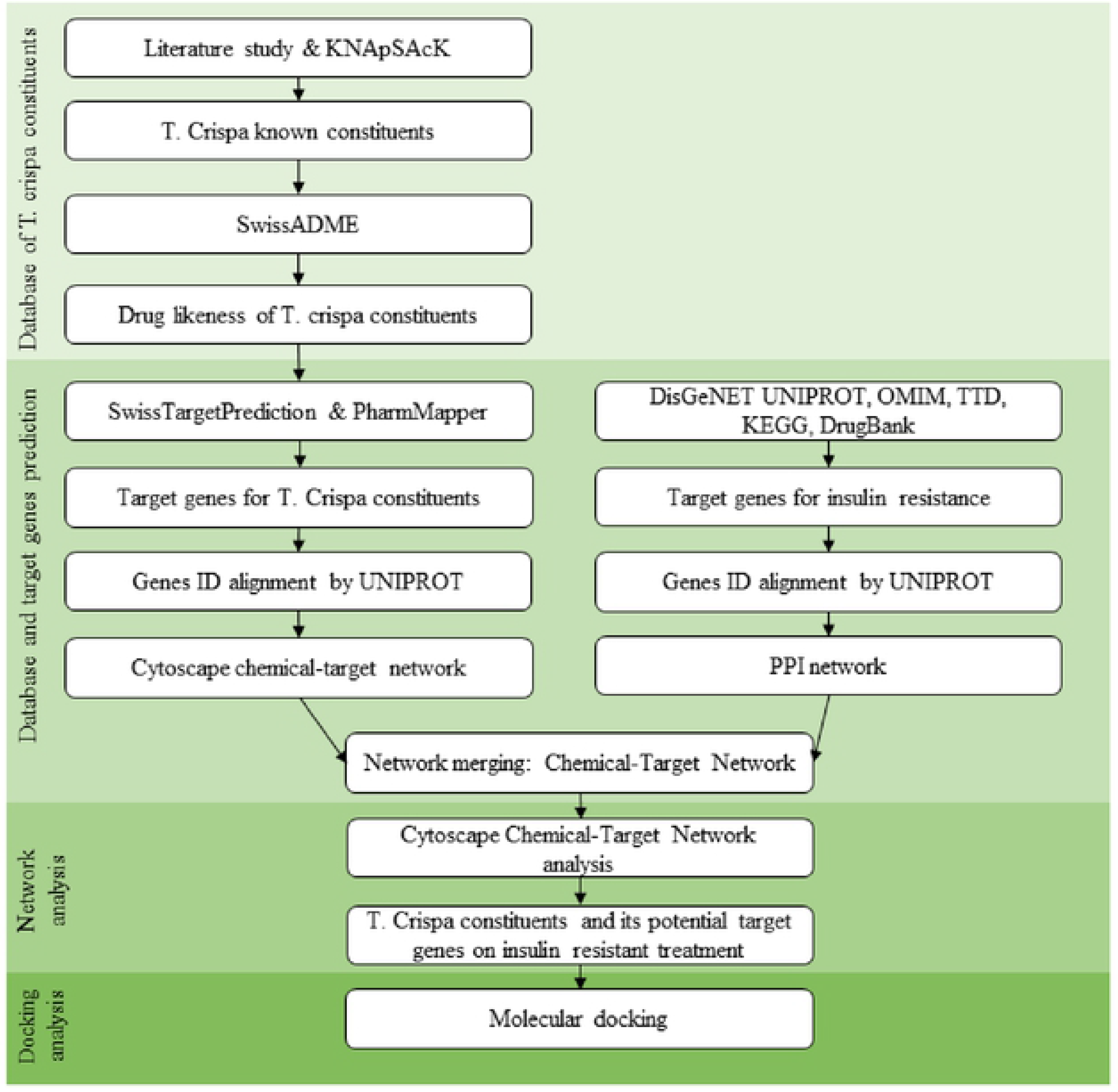
General workflow of present work using network pharmacology and molecular docking analysis.

## Material and Methods

### Constructing database of known *Tinospora crispa* chemical constituents

Data of chemical constituents contained in *T. crispa* was acquired from former researches related to *T. crispa* constituent identification^4–12^. Database constructing of 2D structure was made in sdf format were collected from PubChem, ChemSpider (http://www.chemspider.com/), or drawn using MarvinSketch (https://chemaxon.com/products/marvin)^14^. Each 2D structure then changed to 3D structure using MarvinSketch and saved as pdb format^15^. The 2D structure was needed in druggability analysis in each component, besides the 3D structure was needed for molecular docking analysis. Its compound’s drugability were predicted by SwissADME database (http://www.swissadme.ch/) including its drug likeness based on Lipinsky rules of five and its oral bioavailability prediction^16^.

### Constructing a database of protein target involved in insulin resistantance

Target proteins related to insulin resistance were retrieved from DisGeNET, UniProt, OMIM, TTD, KEGG, and DrugBank database^17–21^. Data retrieval from DisGeNET were done with keyword “Insulin resistance” (CUI: C0021655) in diseases search engine coloumn. The retrieved data was a summary of gene disease association that have been downloaded in xlsx format (80 genes). Data retrieval from UniProt was done using human disease option in supporting data and input keyword “Diabetes mellitus, non-insulin-dependent” (DI-02060). Data were downloaded as tab file format (23 genes). Data from both data sources were merging in Cytoscape 3.7.2 software (https://cytoscape.org/) using UniProt ID as their identity to eliminate double listed proteins^18, 22^.

### Constructing Target Proteins of *Tinospora crispa* phytochemical Constituents

The simplified molecular input line entry specification (SMILES) information of 56 TC constituents were submitted in SWISS Target Prediction (http://www.swisstargetprediction.ch/) and PharmMapper to obtain its target prediction. Only human (*Homo sapiens*) target protein was set in the data retrieval^23, 24^. Protein ID were aligned using UniProt ID to synchronize protein ID and eliminate double listed protein^18^. The aligned target proteins ID were arranged in excel format to be imported to Cytoscape 3.7.2, be visualized, and analyzed its degree of connectivity^22^.

### Prediction of significance target protein related insulin resistant

Protein-protein interaction (PPI) was generated by submitting all UniProt ID of the proteins which have been constructed from the previous step into STRING database (https://string-db.org/) using multiple proteins option^25^. *Homo sapiens* was set as the only organism and the interaction score confidence was set at the highest confidence (0.900). PPI data then downloaded in tsv format. Furthermore, the PPI network was analyzed using Cytoscape 3.7.2 software to discover the rank of significance protein based on the degree of connectivity score^22^.

### Prediction of *Tinospora crispa* phytoconstituents target genes and their intersection on insulin resistant related target

All the phytochemical *T. crispa* compounds were predicted its target by SwissTargetPrediction and PharmMapper^23, 24^. To comprehensively understand its molecular mechanism, all compound-target data were visualized by Cytoscape 3.7.2 to reveal the interaction network between compound-target. Furthermore, the network then merged with PPI network to discover a new network, compound-target network of *T. crispa* targeted to insulin resistance.

### Ligand and protein preparation for in silico molecular docking

Molecular docking was conducted towards the significant proteins in insulin resistance pathogenesis and was targeted by constituents of *T. crispa* at once. Target proteins that meet the criteria were PIK3R1, PTPN1, PPARG, INSR, EGFR, TNF, and AKT2. The 3D structure of *T. crispa* constituents act as ligands (file type .pdb) were opened using AutoDock Tools 1.5.6. software (Molecular Graphics Laboratory, the Scripps Research Institute)^15^. Water molecules were removed and chain contained active site was chosen. The chosen structure then extracted from its native ligand. The 3D crystal structures of target proteins were selected from Protein Data Bank (PDB) (http://www.rcsb.org/pdb/) as shown in Fig 3A and Supplementary material 4^26^. Protein structure with PDB accession number 3S2A were used for PIK3R1 target, 1LQF for PTPN1 target, 6O67 for PPARG target, 5E1S for INSR target, 6D8E for EGFR target, 5MU8 for TNF target, and 5D0E for AKT2 target. All protein models were prepared by the addition of polar hydrogen and the addition of Gasteiger charges. Ligands were prepared with torsion value less than 32. Both ligands and protein models were saved as pdbqt file. Preparation steps were done using AutoDock Tools 1.5.6 software.

### Molecular docking

Molecular docking was generated using AutoDock Vina (Molecular Graphics Laboratory, the Scripps Research Institute) for initial screening of all *T. crispa* constituents followed by AutoDock Tools 1.5.6 only for the top 3 constituents with the lowest binding energy^15, 27^. The ligands were set as rigid structures with active site was set visually at the center of the cavity docked by native ligand from each protein model. The grid box was adjusted at 40 x 40 x 40 Angstrom in x, y, and z axis respectively. Molecular docking was run in Lamarckian Genetic Algorithm after validation. Redocking of native ligand with its protein in the certain parameters with RMSD value less than 2 Amstrong remains as a valid docking parameter^28^. The results were shown in value of its lowest binding energy (kcal/mol) calculated by total intermolecular energies including hydrogen bonds energy, Van der Walls energy, desolvation energy, and electrostatic energy. The more negative energy was, the better ligand-protein binding^29^. The result of ligan-protein complexes were visualized using PyMOL (https://pymol.org/2/). The docking results were analyzed in LigPlot+ to evaluate the accurate binding interaction between ligands and proteins^30^.

## Results

### Database of *Tinospora crispa* phytoconstituents

A total of 56 compounds were found in literatures related to *T.crispa*. 2D and 3D structures of all compounds were collected from PubChem, ChemSpider, or drawn using Marvin Sketch^14^. Its compounds druggability were predicted using SwissADME including its drug likeness based on Lipinsky rules of five and its oral bioavailability prediction (Table 1)^16^. The molecular weight range from 135.13 to 714.71 with Adenine as the lightest molecule and Borapetoside H as the heaviest molecule. Predicted gastrointestinal absorption showed 34 of 56 compounds classified as highly absorbed in the GI tract. Drug likeness was assessed through the Lipinski Rules of Five, with 37 compounds meet the 5 criteria required by Lipinski, while 19 other compounds have violations of Lipinski’s rules of 1-3 violations.

**Table 1.**
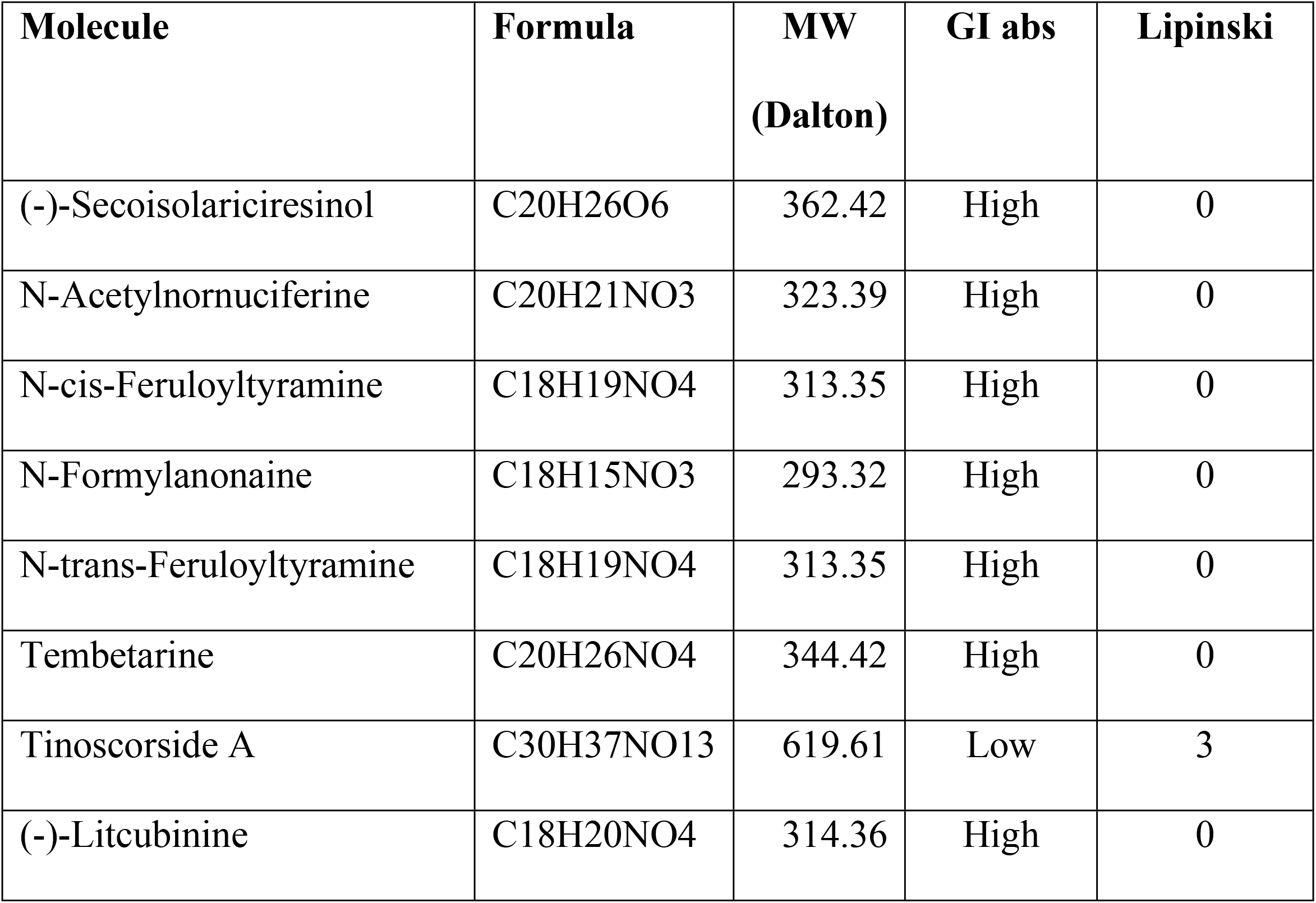
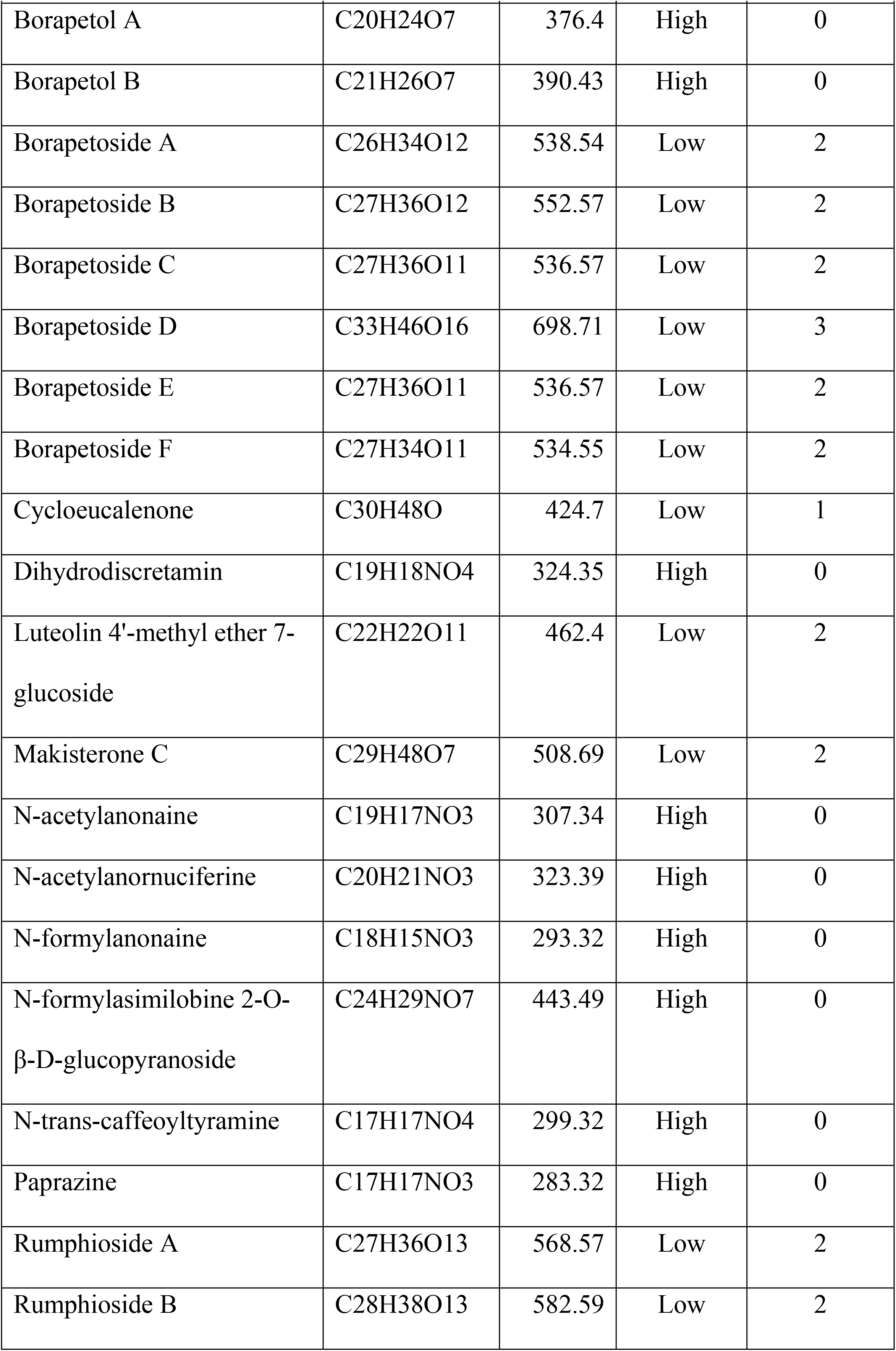
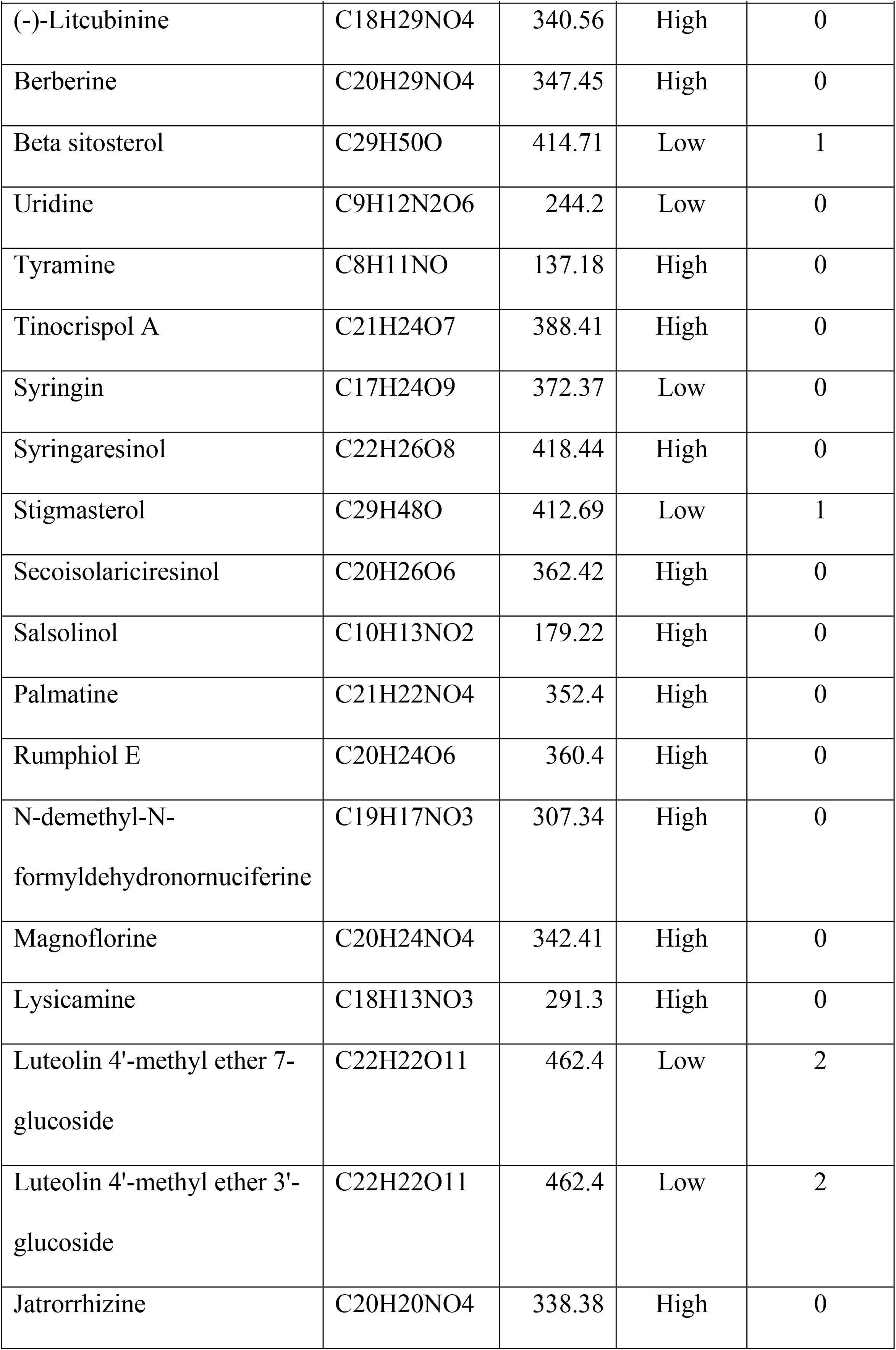
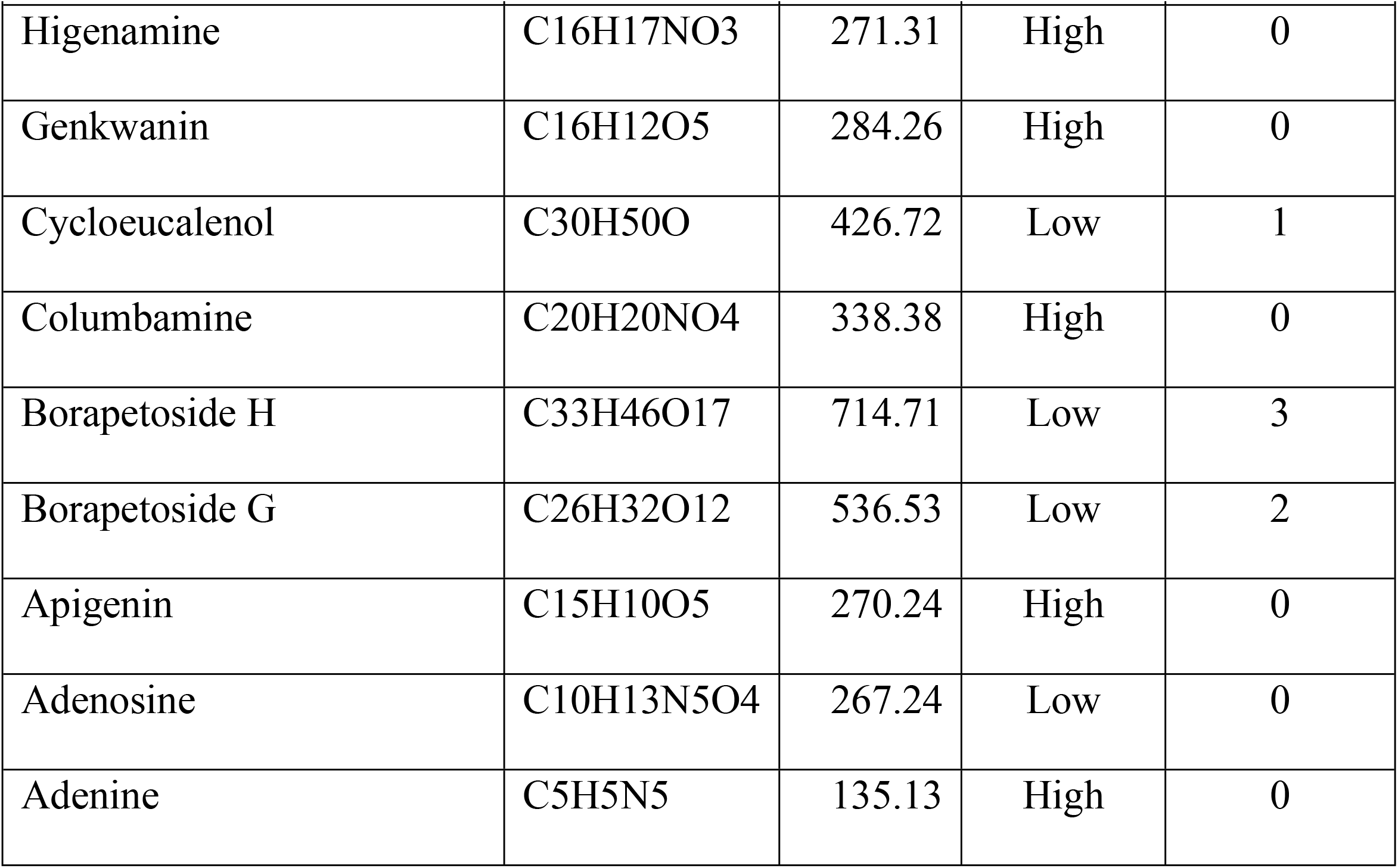
Prediction of *Tinospora crispa* know phytoconstituents and its druggability^16^

| Molecule | Formula | MW<br>(Dalton) | GI abs | Lipinski |
| --- | --- | --- | --- | --- |
| (-)-Secoisolariciresinol | C<sub>20</sub>H<sub>26</sub>O<sub>6</sub> | 362.42 | High | 0 |
| N-Acetylnornuciferine | C<sub>20</sub>H<sub>21</sub>NO<sub>3</sub> | 323.39 | High | 0 |
| N-cis-Feruloyltyramine | C<sub>18</sub>H<sub>19</sub>NO<sub>4</sub> | 313.35 | High | 0 |
| N-Formylanonaine | C<sub>18</sub>H<sub>15</sub>NO<sub>3</sub> | 293.32 | High | 0 |
| N-trans-Feruloyltyramine | C<sub>18</sub>H<sub>19</sub>NO<sub>4</sub> | 313.35 | High | 0 |
| Tembetarine | C<sub>20</sub>H<sub>26</sub>NO<sub>4</sub> | 344.42 | High | 0 |
| Tinoscorside A | C<sub>30</sub>H<sub>37</sub>NO<sub>13</sub> | 619.61 | Low | 3 |
| (-)-Litcubinine | C<sub>18</sub>H<sub>20</sub>NO<sub>4</sub> | 314.36 | High | 0 |

| **Molecule** | **Formula** | **MW**<br>**(Dalton)** | **GI abs** | **Lipinski** |
| --- | --- | --- | --- | --- |
| Borapetol A | C20H24O7 | 376.4 | High | 0 |
| Borapetol B | C21H26O7 | 390.43 | High | 0 |
| Borapetoside A | C26H34O12 | 538.54 | Low | 2 |
| Borapetoside B | C27H36O12 | 552.57 | Low | 2 |
| Borapetoside C | C27H36O11 | 536.57 | Low | 2 |
| Borapetoside D | C33H46O16 | 698.71 | Low | 3 |
| Borapetoside E | C27H36O11 | 536.57 | Low | 2 |
| Borapetoside F | C27H34O11 | 534.55 | Low | 2 |
| Cycloeucalenone | C30H48O | 424.7 | Low | 1 |
| Dihydrodiscretamin | C19H18NO4 | 324.35 | High | 0 |
| Luteolin 4'-methyl ether 7-glucoside | C22H22O11 | 462.4 | Low | 2 |
| Makisterone C | C29H48O7 | 508.69 | Low | 2 |
| N-acetylanonaine | C19H17NO3 | 307.34 | High | 0 |
| N-acetylanornuciferine | C20H21NO3 | 323.39 | High | 0 |
| N-formylanonaine | C18H15NO3 | 293.32 | High | 0 |
| N-formylasimilobine 2-O-β-D-glucopyranoside | C24H29NO7 | 443.49 | High | 0 |
| N-trans-caffeoyltyramine | C17H17NO4 | 299.32 | High | 0 |
| Paprazine | C17H17NO3 | 283.32 | High | 0 |
| Rumphioside A | C27H36O13 | 568.57 | Low | 2 |
| Rumphioside B | C28H38O13 | 582.59 | Low | 2 |
| (-)-Litchubinine | C18H29NO4 | 340.56 | High | 0 |
| Berberine | C20H29NO4 | 347.45 | High | 0 |
| Beta sitosterol | C29H50O | 414.71 | Low | 1 |
| Uridine | C9H12N2O6 | 244.2 | Low | 0 |
| Tyramine | C8H11NO | 137.18 | High | 0 |
| Tinocrispol A | C21H24O7 | 388.41 | High | 0 |
| Syringin | C17H24O9 | 372.37 | Low | 0 |
| Syringaresinol | C22H26O8 | 418.44 | High | 0 |
| Stigmasterol | C29H48O | 412.69 | Low | 1 |
| Secoisolariciresinol | C20H26O6 | 362.42 | High | 0 |
| Salsolinol | C10H13NO2 | 179.22 | High | 0 |
| Palmatine | C21H22NO4 | 352.4 | High | 0 |
| Rumphiol E | C20H24O6 | 360.4 | High | 0 |
| N-demethyl-N-formyldehydronornuciferine | C19H17NO3 | 307.34 | High | 0 |
| Magnoflorine | C20H24NO4 | 342.41 | High | 0 |
| Lysicamine | C18H13NO3 | 291.3 | High | 0 |
| Luteolin 4'-methyl ether 7-glucoside | C22H22O11 | 462.4 | Low | 2 |
| Luteolin 4'-methyl ether 3'-glucoside | C22H22O11 | 462.4 | Low | 2 |
| Jatrorrhizine | C20H20NO4 | 338.38 | High | 0 |
| Higenamine | C16H17NO3 | 271.31 | High | 0 |
| Genkwanin | C16H12O5 | 284.26 | High | 0 |
| Cycloeucalenol | C30H50O | 426.72 | Low | 1 |
| Columbamine | C20H20NO4 | 338.38 | High | 0 |
| Borapetoside H | C33H46O17 | 714.71 | Low | 3 |
| Borapetoside G | C26H32O12 | 536.53 | Low | 2 |
| Apigenin | C15H10O5 | 270.24 | High | 0 |
| Adenosine | C10H13N5O4 | 267.24 | Low | 0 |
| Adenine | C5H5N5 | 135.13 | High | 0 |

From SWISS target prediction and DisGeNET, identified that 56 constituents of *T. crispa* were documented to have 5666 protein-specific targets to insulin resistance^17, 23^. Those 5666 target proteins then analyzed by Cytoscape 3.7.2 to find out its intersection with insulin resistant target protein^22^.

### Protein-protein interaction networks related to insulin resistance

267 targets were documented to have relation to insulin resistance pathogenesis. All targets were identified its interaction using STRING database^25^. Those interactions construct a PPI network which then assessed by Cytoscape 3.7.2 software for the degree of connectivity value. The analysis results were displayed in Fig 2 in white to purple gradation. The higher the degree of a node (gene) in the network is marked with a darker purple color. The higher the degree value indicates that the protein has a greater role in the pathogenesis of insulin resistance. The top degree ranks were INS, LEP, PIK3R1, PTPN1, IRS1, PPARG, IGF1, INSR, CAV1, EGFR, IRS2, TNF, and AKT2, respectively. The degree rating of the entire PPI network is shown in Fig 2. The detailed degree value was shown in Supplementary material 1.

**Fig 2.**
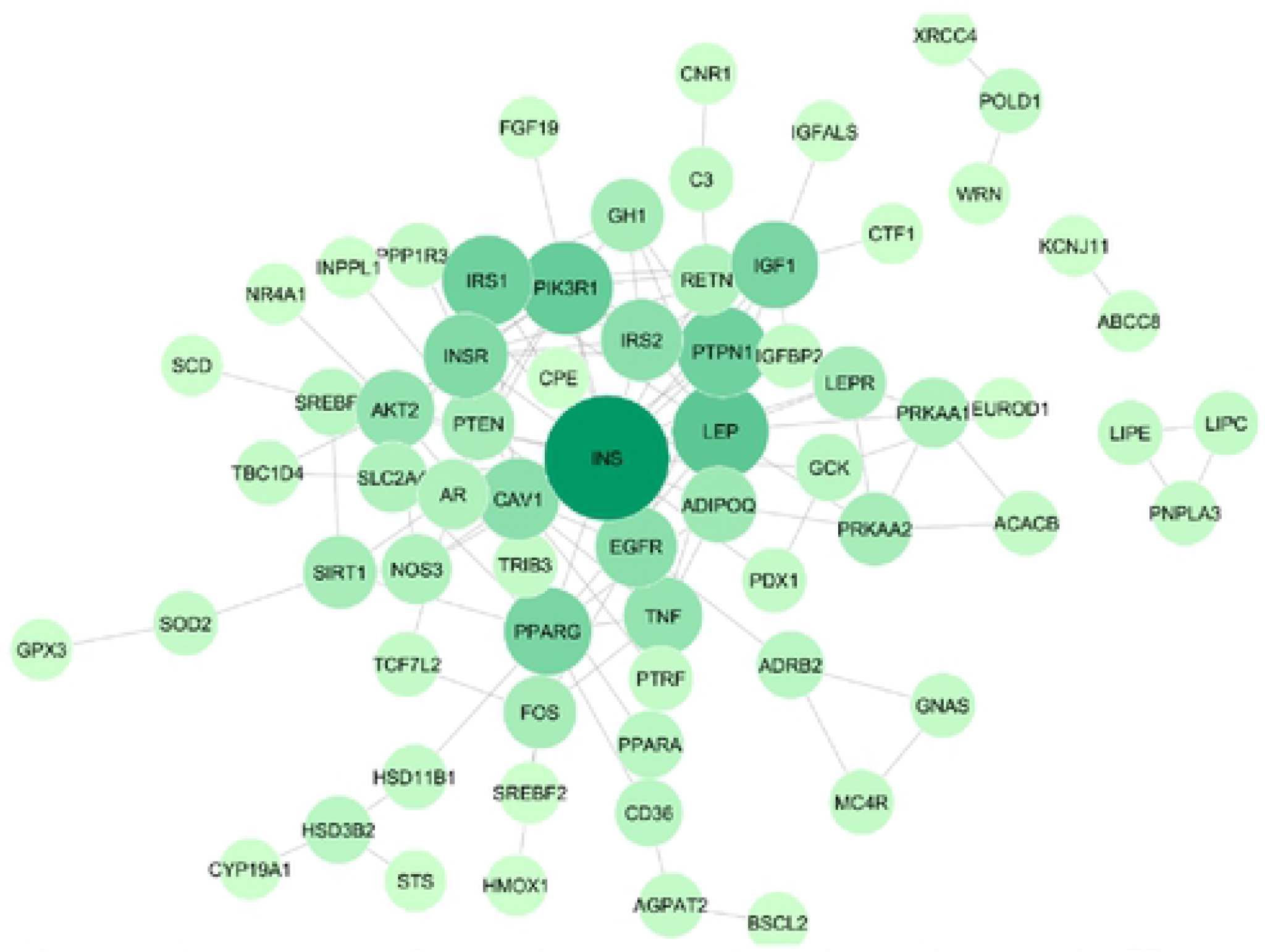
PPI network of protein related insulin resistance in *Homo sapiens.* The darker purple color and the bigger its nodes indicate higher degree of protein in the network.

### Interaction of lead compounds of *Tinospora crispa* on insulin resistant related target proteins

To proof the underlying interactions between *T. crispa* predicted active phytoconstituents to insulin resistance, we conducted the molecular docking strategy. Those targets were PIK3R, PTPN1, PPARG, INSR, EGFR, TNF, and AKT2. The best docking interaction was PI3K with Tinoscorside A as a ligand (binding energy -11.64) as shown in Table 2. Its binding sketch map was shown in Fig 3 as well. The detail docking scores of *T. crispa* targeted to 7 significance proteins were shown in Supplementary material 3 and the docking sketch maps were shown in Supplementary material 4. The structures of main constituents of T. *crispa* were showed in Fig 4. Seven target proteins that meet the criteria to have a high degree of connectivity value, targeted by *T. crispa* constituents, and were involved in insulin resistance pathogenesis (Fig 5) were used in this docking analysis.

**Table 2.** Docking scores of the lowest ligand-protein binding energy of T. crispa constituents and significant proteins in insulin resistance.

| Target protein | Compound | Ki (nM) | The lowest binding energy (kcal/mol) |
| --- | --- | --- | --- |
| PI3K | Tinoscorside A | 2.94 | -11.64 |
| INSR | Tinoscorside A | 4.19 | -11.43 |
| AKT2 | Makisterone C | 5.17 | -11.3 |
| PPARG | Borapetoside A | 51.96 | -9.94 |
| EGFR | Tinoscorside A | 191.99 | -9.16 |
| PTPN1 | Beta sitosterol | 453.59 | -8.65 |
| TNF | Borapetoside B | 1.62x10<sup>3</sup> | -7.90 |

**Fig 3.**
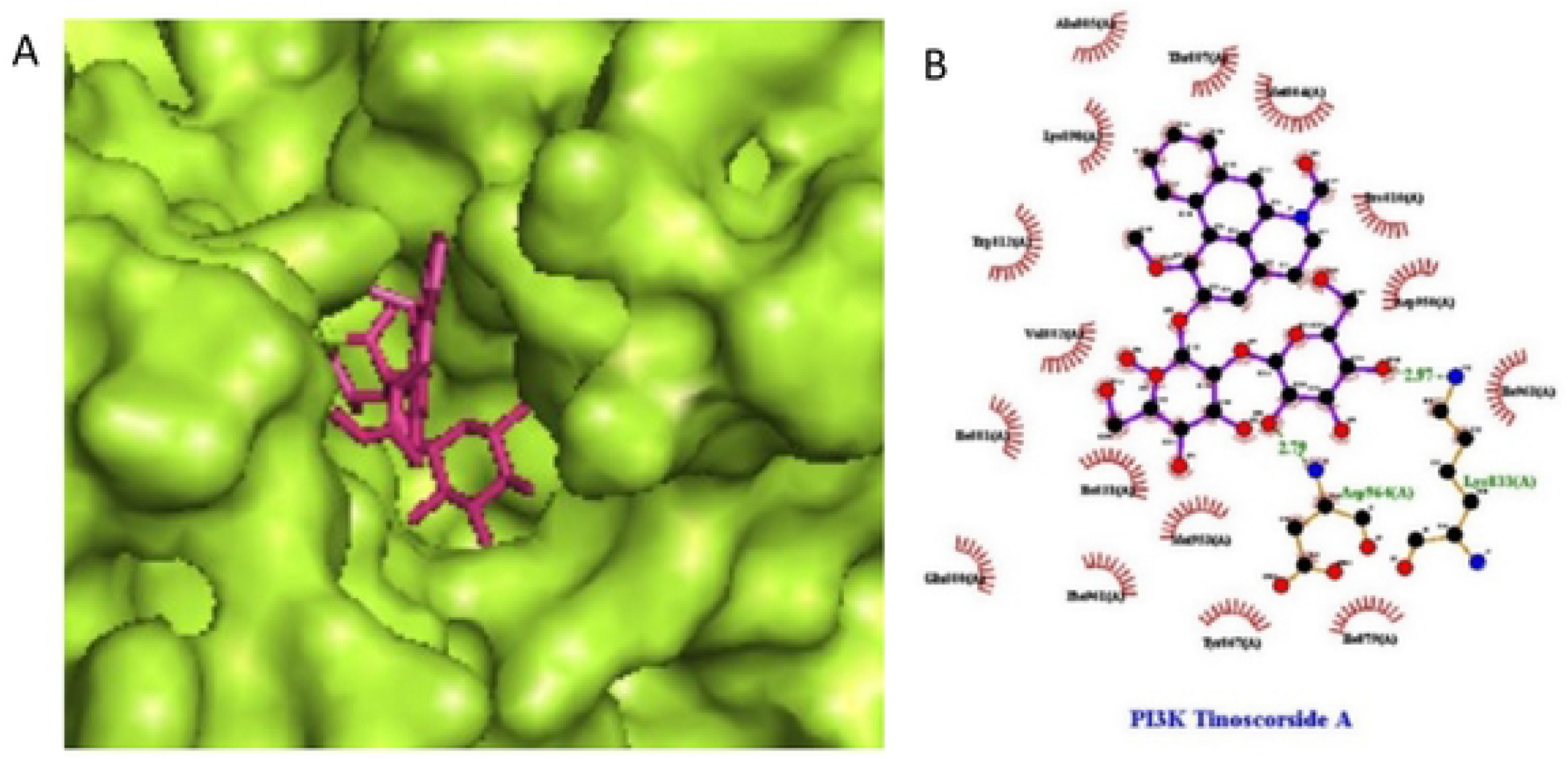
Docking sketch map of Tinoscorside A interaction with the crystal structure of PI3K. (A) Pocket view of Tinoscorside A binding with PI3K active site. (B) 2D docking pattern of Tinoscorside A with amino acids hydrogen bonds LYS833 and ASP964.

**Fig 4.**
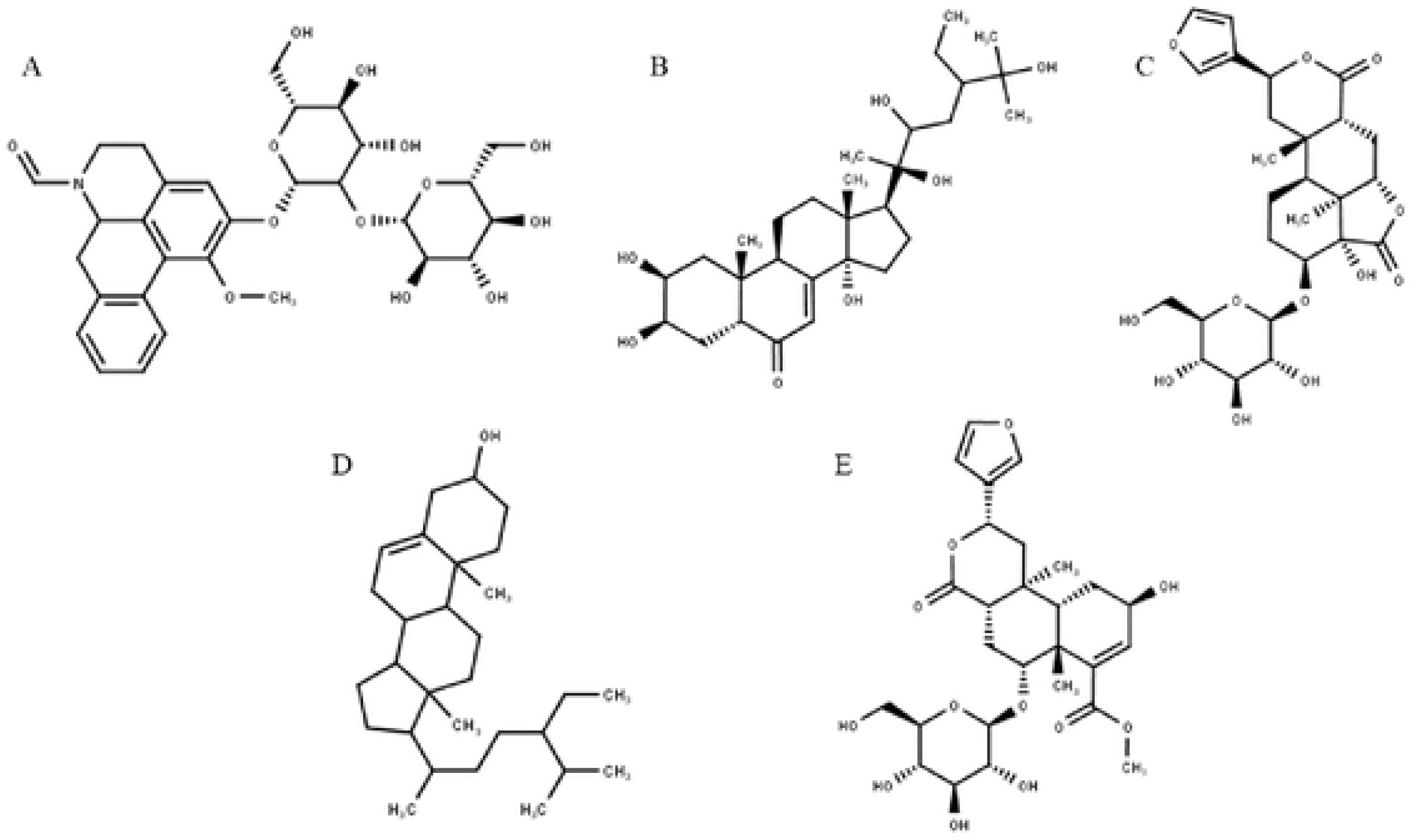
Structure of active constituents prediction of *T. crispa* as insulin sensitizer. (A) Tinoscorside A (B) Makisterone C (C) Borapetoside A (D) β Sitosterol (E) Borapetoside B. All structures were drawn using MarvinSketch based on the previous studies^6^.

**Fig 5.**
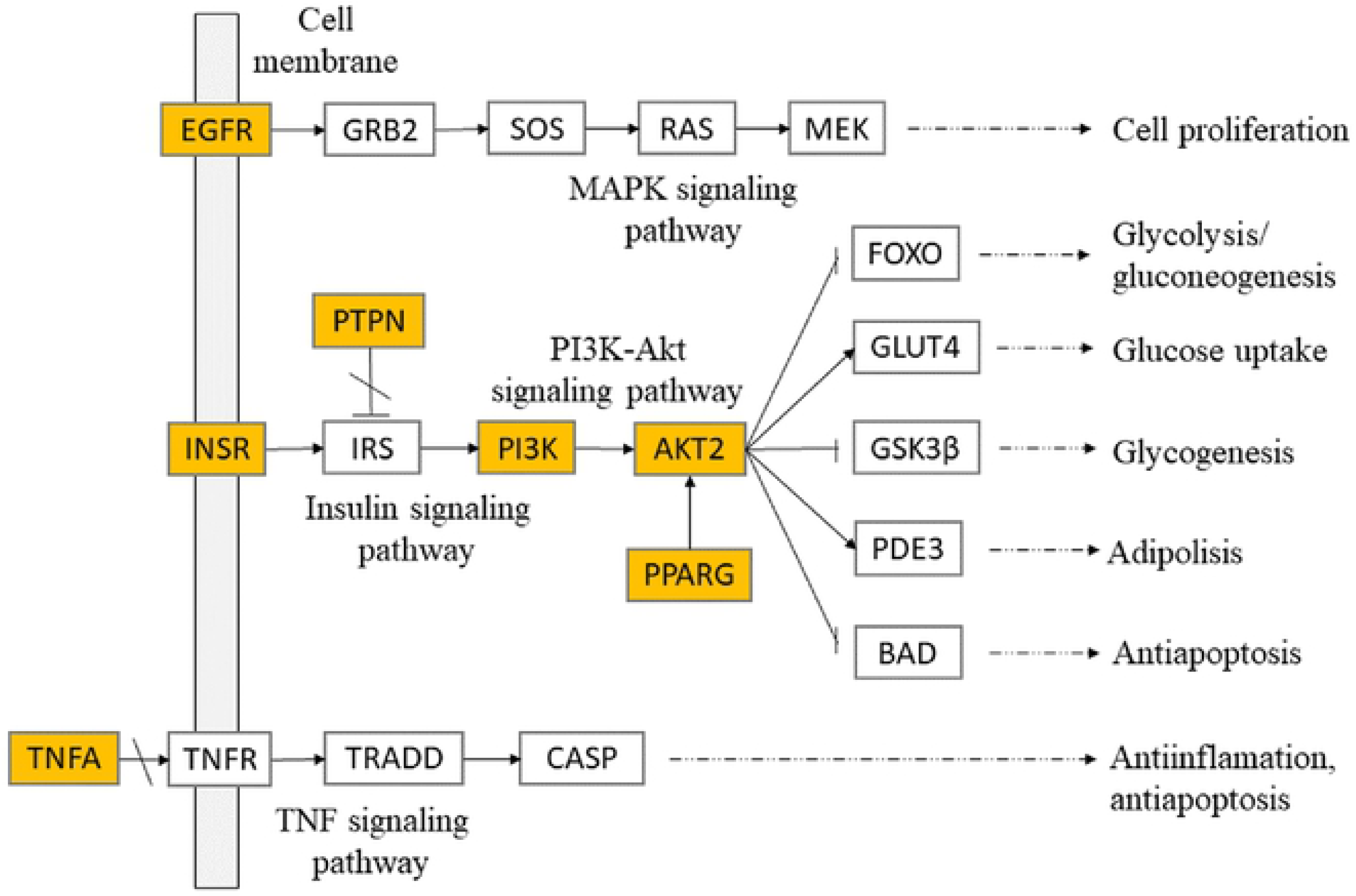
Systematic prediction of 7 significant protein targets and therapeutic pathways of *T. crispa* constituents on insulin resistant. The highlighted nodes represent the significant targets of *T. crispa* and other nodes represent their related target on insulin resistant. All the presented pathways were summarized by KEGG pathway database^21^.

### Compound-Target Network

To further discover the potential target and therapeutic pathway of *T. crispa* constituents in ameliorating insulin resistance, the targets of insulin resistance were merged with the target of *T. crispa* (Fig 6A). The compound-target network of *T. crispa* against insulin resistance was shown in Fig 6C. After the network merging using Cytoscape 3.7.2, 30 *T. crispa* constituents were reserved as the main active constituents in *T. crispa* treating insulin resistance. In addition, 22 targets were reserved as the main targets of TC in treating insulin resistance. Moreover, 7 targets were revealed to have significant protein in insulin resistance pathogenesis and targeted by *T. crispa* constituents (Fig 6B). The complete *T. crispa* constituents and their targets were shown in Supplementary material 2.

**Fig 6.**
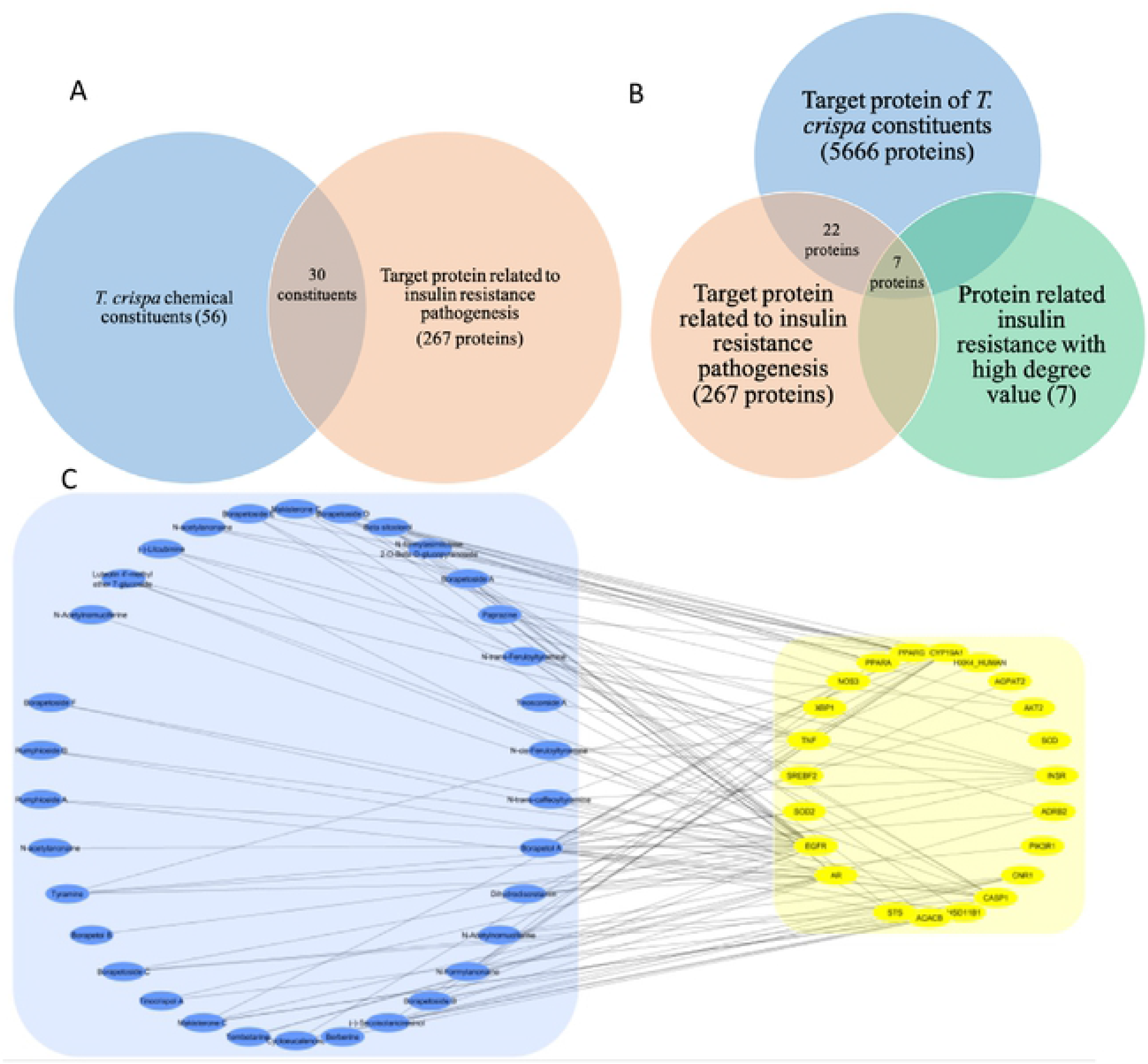
**(A) Intersection Venn diagram of *T. crispa* constituents and target protein related insulin resistance.** There were 30 constituents of *T. crispa* that were predicted to be targeted to insulin resistance target protein. **(B) Intersection Venn diagram between predicted target protein of *T. crispa* constituents, target proteins of insulin resistance pathogenesis, and high degree of connection value of target proteins of insulin resistance.** There were 7 target proteins that meet the three intersections. In addition there were 22 proteins targeted by *T. crispa* constituents that were shown visually in figure 6C. **(C) Compound-target network of *T. crispa* constituents targeted to insulin resistance pathogenesis.** The blue dots represent *T. crispa* constituents, the yellow dots represent protein targets, and the connecting lines represent compound-target interaction.

## Discussion

The present study tried to collect and investigate clues from the known phytoconstituents of *T. crispa* in ameliorating insulin resistance. Investigations were conducted using *in silico* approaches including network pharmacology and molecular docking. This analysis informed that the vital phytoconstituents of *T. crispa* as insulin sensitizer were Tinoscorside A, Makisterone C, Borapetoside A, Beta sitosterol, and Borapetoside B. In addition, we elucidated the potential therapeutic network of insulin resistance and intersect this network to the *T. crispa* phytoconstituents potential targets network. Both networks were complicated networks with 5666 and 267 nodes. The intersection showed us only the protein related insulin resistance targeted by *T. crispa* constituents. From 267 targets, there were only 22 target proteins that targeted by *T. crispa* constituents. KEGG pathways analysis had indicated several signaling pathways including insulin signaling, PI3K-Akt signaling, MAPK signaling, and TNF signaling pathway that play vital roles in the therapeutic mechanism of *T. crispa* in ameliorating insulin resistance. Those signaling pathways had close relative to glucose lowering activity through the regulation of glucose homeostasis, adipolysis, cell proliferation, and antiapoptosis. This showed that *T. crispa* with its various secondary metabolites had activity as an insulin sensitizer through different pathways. This finding supports Wink *et al* statement that the activity of a medicinal plant was due to synergistic interactions of several phytoconstituents.^13^. The network pharmacology approach provides an insight into prediction in the mode of action of a medicinal plant in a comprehensive manner with multiple target therapies^31^.

Based on former studies, Borapetoside A-C, and E may be the key of *T. crispa* constituents as hypoglycaemic agents^7, 11, 12, 32, 33^. Lam *et al* reported, the isolated Borapetoside A and C had significantly reduced plasma glucose levels but unable to increase insulin secretion in MIN6 pancreatic beta cells^32^. It indicated that they had other mechanisms of hypoglycemic activity. This study predicts the mechanism of action of Borapetoside A through activation of PPARG and Borapetoside B through inhibition of TNF cytokines to bind to their receptor.

Contrary to previous studies that indicated diterpenoids as active constituents, this study presents Tinoscorside A (N-formylasimilobine 2-O-β-D-glucopyranosyl-(1→2)-β-D-glucopyranoside) as aconstituent of *T. crispa* which has the strongest binding with some protein targets including PI3K, INSR, and EGFR respectively. Tinoscorsida A was a member of aporphine alkaloid which has never been proven as part of active compounds of T. crispa in reducing blood sugar level. This finding indicated that Tinoscorside A could be a potential glucose lowering agent through multiple ways. In addition, this study also predicted that T. crispa had anti-inflammatory activity through the TNF signaling pathway by inhibit the binding between TNFA and TNFR.

## Conclusion

The present study highlights the insulin sensitizing role of *T. crispa* on insulin resistance pathogenesis by network pharmacology and molecular docking study. The role of *T. crispa* phytoconstituents that served as plant extract with multiple constituents remains still undescribed. Our study provides functional clues to reveal the prediction of their possible targets and mechanism pathways of *T. crispa.* The result was showed that *T. crispa* was a promising plant-based drug to be developed as insulin sensitizer through multiple pathways including glucose homeostasis, adipolysis, cell proliferation, and antiapoptosis. However, there are several limitations in the current study, including the limitation of *T. crispa* phytoconstituents database that could be more phytoconstituents were not reveal yet and the limitation of the database that serves the activity of the constituents as a single compound neglecting the interaction of the multi compound itself. Further study on the insulin sensitizing role of *T. crispa* and the underlying mechanism are still urgently to be investigated in other methods such as *in vitro*, *in vivo,* and clinical study evaluation to verify this *in silico* study.

## Acknowledgments

This study was funded by Universitas Indonesia research grants for doctoral students through PUTI Grant with contract number NKB585/UN2.RST/HKP05.00/2020.

## Supporting Information

### S1

**Table S1.** Degree ranking of insulin resistant related targets analyzed by Cytoscape 3.7.2.

| Gene<br>name | Uniprot<br>ID | Degree |  | Gene<br>name | Uniprot<br>ID | Degree |
| --- | --- | --- | --- | --- | --- | --- |
| INS | P01308 | 23 |  | MC4R | P32245 | 2 |
| LEP | P41159 | 13 |  | ACACB | O00763 | 2 |
| PIK3R1 | P27986 | 12 |  | HSD11B1 | P28845 | 2 |
| PTPN1 | P18031 | 11 |  | TCF7L2 | Q9NQB0 | 2 |
| IRS1 | P35568 | 11 |  | TRIB3 | Q96RU7 | 2 |
| PPARG | P37231 | 10 |  | LIPC | P11150 | 2 |
| IGF1 | P05019 | 10 |  | POLD1 | P28340 | 2 |
| INSR | P06213 | 9 |  | GNAS | Q5JWF2 | 2 |
| CAV1 | Q03135 | 8 |  | PTRF | Q6NZI2 | 2 |
| EGFR | P00533 | 8 |  | SOD2 | P04179 | 2 |
| IRS2 | Q9Y4H2 | 8 |  | PNPLA3 | Q9NST1 | 2 |
| TNF | P01375 | 7 |  | LIPE | Q05469 | 2 |
| AKT2 | P31751 | 7 |  | PDX1 | P52945 | 2 |
| ADIPOQ | Q15848 | 6 |  | TBC1D4 | O60343 | 2 |
| GH1 | P01241 | 5 |  | IGFBP2 | P18065 | 2 |
| PRKAA2 | P54646 | 5 |  | FGF19 | O95750 | 1 |
| SIRT1 | Q96EB6 | 5 |  | NR4A1 | P22736 | 1 |
| FOS | P01100 | 5 |  | CNR1 | P21554 | 1 |
| PTEN | P60484 | 5 |  | STS | P08842 | 1 |
| LEPR | P48357 | 5 |  | XRCC4 | Q13426 | 1 |
| PRKAA1 | Q13131 | 4 |  | BSCL2 | Q96G97 | 1 |
| RETN | Q9HD89 | 4 |  | CTF1 | Q16619 | 1 |
| AR | P10275 | 4 |  | SCD | O00767 | 1 |
| SLC2A4 | P14672 | 4 |  | HMOX1 | P09601 | 1 |
| NOS3 | P29474 | 4 |  | SREBF2 | Q12772 | 1 |
| SREBF1 | P36956 | 3 |  | INPPL1 | O15357 | 1 |
| PPARA | Q07869 | 3 |  | NEUROD1 | Q13562 | 1 |
| CD36 | P16671 | 3 |  | WRN | Q14191 | 1 |
| ADRB2 | P07550 | 3 |  | CPE | P16870 | 1 |
| GCK | P35557 | 3 |  | CYP19A1 | P11511 | 1 |
| HSD3B2 | P26439 | 3 |  | GPX3 | P22352 | 1 |
| C3 | P01024 | 2 |  | IGFALS | P35858 | 1 |
| AGPAT2 | O15120 | 2 |  | ABCC8 | Q09428 | 1 |
| PPP1R3A | Q16821 | 2 |  | KCNJ11 | Q14654 | 1 |

### S2

**Table S2.**
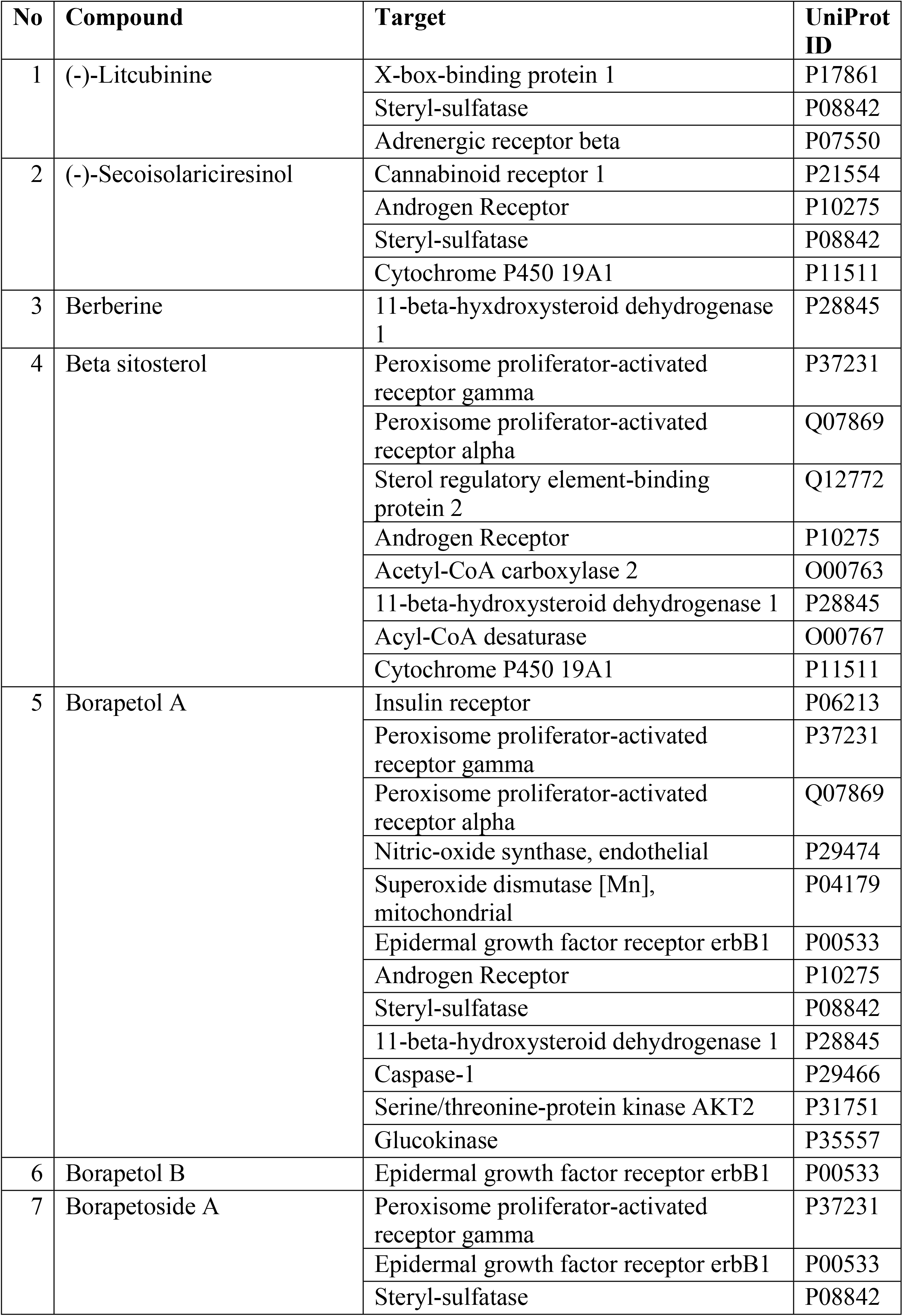
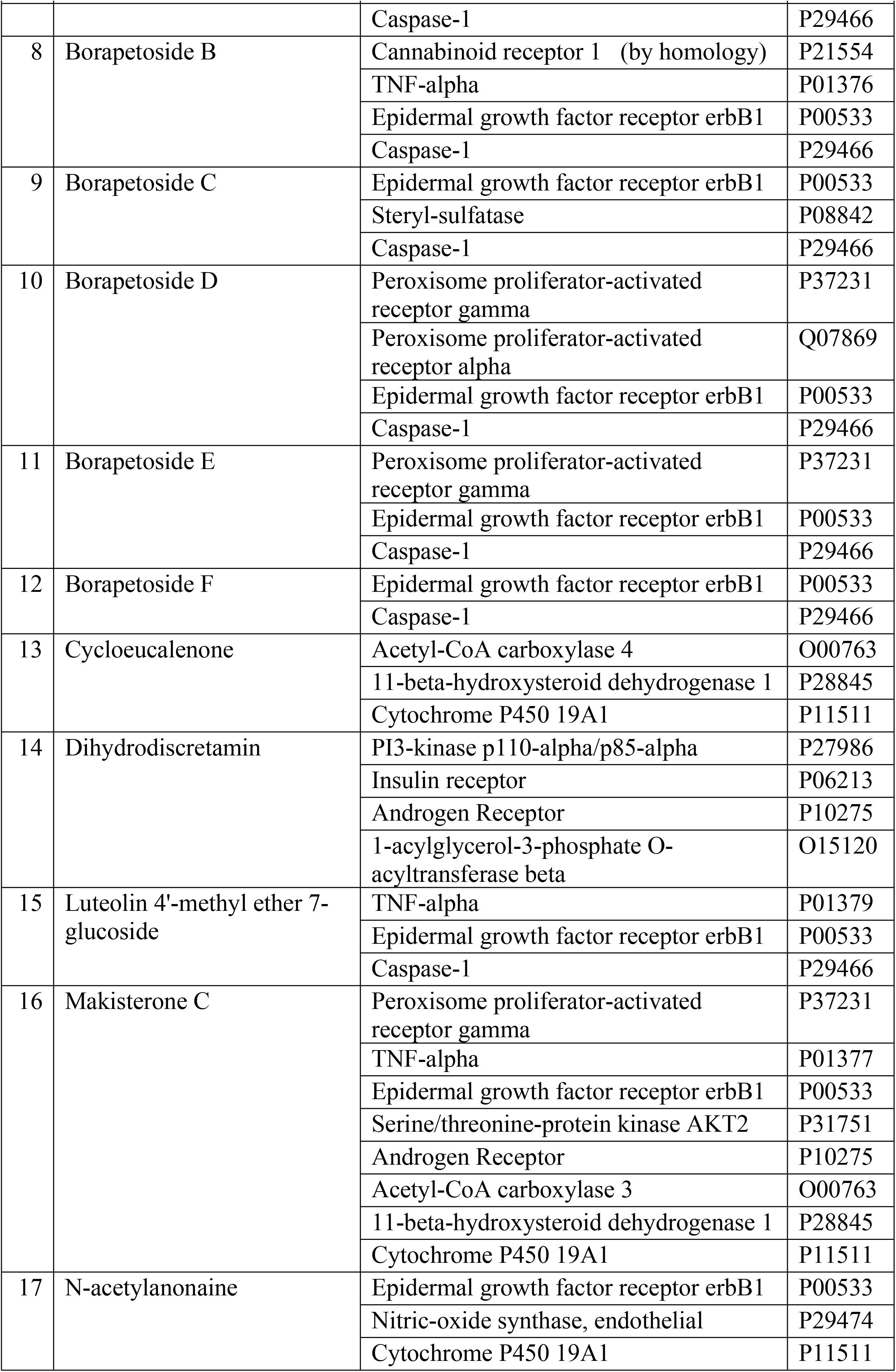
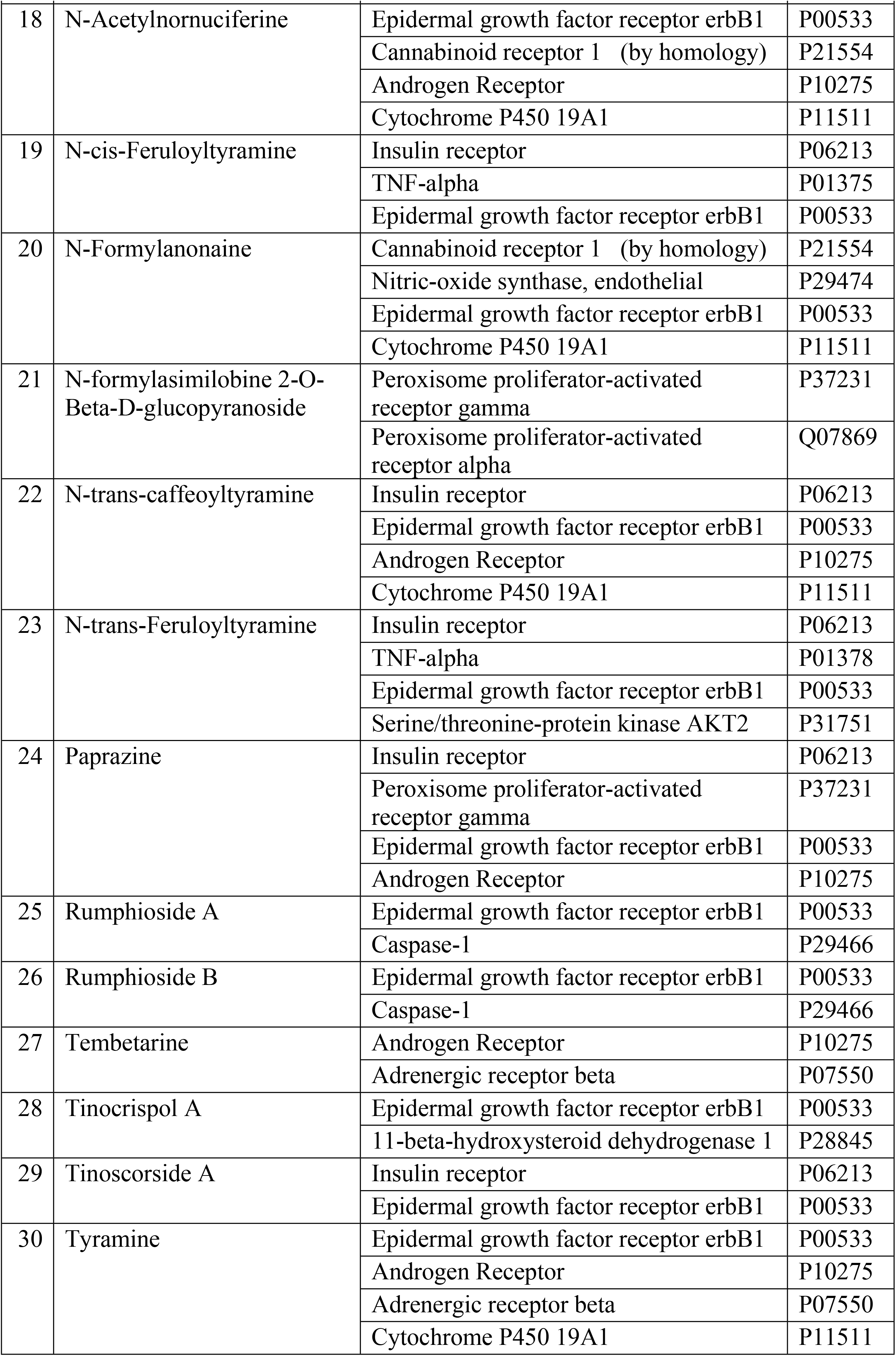
30 T. crispa phytoconstituents and its target proteins^17, 18, 24^.

### S3

**Table S3.** Molecular docking result of *T. crispa* phytoconstituents to 7 selected target proteins using AutoDock.

| Protein target | TC Compound | Lowest binding energy | Hydrogen bond | Hydrophobic bond |
| --- | --- | --- | --- | --- |
| PI3K | Tinoscorside A | **-11.64** | Asp96, Lys833 | Ala 805, Thr 887, Met 804, Pro 810, Asp 950. Ile 953, Ile 879, Tyr 867, Met 953, Ile 831, phe 961, Glu 880, Ile 881, Val 882, Trp 812, Lys 890 |
| PI3K | Borapetoside A | -9.88 |  | Asn 951, Ile 963, Lys 833, Lys 807, Asp 964, Ile 831, Val 882, Tyr 867, Glu 880, Ile 879, Ile 881, Met 804, Ser 806, Pro 810 |
| PI3K | *N-formylasimilobine 2-O-Beta-D-glucopyranoside* | -9.65 | Val 883, Lys 890 | Phe 961, Glu 880, Ile 881, Ala 885, Met 953, Ile 879, Tyr 867, Ile 831, Ile 963, Asn 951, Asp 950, Thr 887, Trp 812 |
| PTPN1 | Beta sitosterol | -8.65 |  | Gly 720, Asp 691, Arg 721, Cys 715, Ala 717, Tyr 546, Phe 482, Asp 548, Ser 716, Gln 762, Ile 719, Gly 759, Val 549. |
| PTPN1 | Jatrorrhizine | -7.79 | Arg 721 | Asp 681, Lys 620, Tyr 546, Ser 716, Asp 548, Val 549, Ile 719, Gln 762, Ala 717, Phe 682. |
| PTPN1 | Magnoflorine | -7.71 | Gly 720, Cys 715, Ala 717, Ser 716, Tyr 546, Lys 620. | Arg 721, Ile 719, Phe 682, Asp 548, Gln 762, Asp 681. |
| PPARG | Borapetoside A | -9.94 |  | Arg 280, Ile 281, Met 348, Cys 285, Leu 353, Leu 330, Ser 342, Leu 333, Ile 341, Glu 343, Leu 340, Gly 284, Arg 288, |
| PPARG | Stigmasterol | -9.71 | Arg 280 | Leu 255, Leu 330, Leu 333, Ile 341, Val 339, Met 348, Leu 340, Ser 342, Arg 288, Cys 285, Gly 284, Glu 343, Phe 287, Ile 281. |
| PPARG | Beta sitosterol | -9.65 |  | Lys 263, Ser 342, Phe 287, Gly 284, Arg 288, Cys 285, Ile 281, Leu 353, Met 364, Met 348, Ile 341, Glu 259, Ile 262, Gly 258. |
| INSR | Tinoscorside A | **-11.43** | Asp 1083, Ser 1086, Met 1079. | His 1081, Gly 1003, Ala 1028, Leu 1078, Val 1010, Leu 1002, Met 1139, Glu 1012, Arg 1000, Arg 1026, Gly 1082, Ala 1080. |
| INSR | N-trans-Feruloyltyramine | -8.85 | Ser 1086, Asp 1083, Met 1079. | Ser 1090, His 1081, Leu 1002, Met 1139, Leu 1078, Gly 1082, Tyr 1087. |
| INSR | N-cis-feruloyltyramine | -8.52 | Met 1079, Asp 1083. | Leu 1078, Ala 1080, Arg 1000, Leu 1002, Ser 1086, Tyr 1087, Ser 1090, His 1081, Met 1139, Gly 1082. |
| EGFR | Tinoscorside A | -9.65 | Asp 855, Thr 845, Cys 797. | Leu 792, Pro 794, Leu 844, Met 793, Ala 743, Thr 790, Met 766, Lys 745, Val 726, Asp 800, Leu 718, Gly 796. |
| EGFR | Borapetoside B | -9.48 |  | Pro 794, Leu 718, Met 793, Leu 844, Thr 790, Ala 743, Thr 854, Asp 855, Asn 842, Arg 841, Cys 797, Leu 792, Gly 796. |
| EGFR | Borapetosida A | -9.21 | Thr 854 | Leu 718, Leu 844, Val 726, Met 793, Ala 743, Thr 790, Leu 792, Ile 789, Leu 777, Leu 788, Met 766, Lys 745, Lys 755, Gly 796. |
| TNF | Borapetoside B | -7.9 |  | Ile 155, Leu 57, Gly 121, Leu 120, Tyr 119, Tyr 151, Tyr 59. |
| TNF | Cycloeucalenol | -6.95 | Gln 149 | Tyr 119, Gln 61, Leu 120, Gly 122, Gly 121, Val 123, Leu 57, Tyr 59, Tyr 151. |
| TNF | Borapetoside H | -5.84 |  | Tyr 151, Tyr 59, Tyr 119, Leu 120, Gly 121, Ser 60. |
| AKT2 | Makisterone C | **-11.3** | Tyr 327, Cys 311, Lys 298, Arg 274, His 196. | - |
| AKT2 | Higenamine | -8.63 | Tyr 327, Cys 311, Lys 298, Arg 274, His 196. | - |
| AKT2 | Borapetol A | -8.17 | Tyr 327, Cys 311, Lys 298, Arg 274, His 196. | - |

### S4

**Fig S4.**
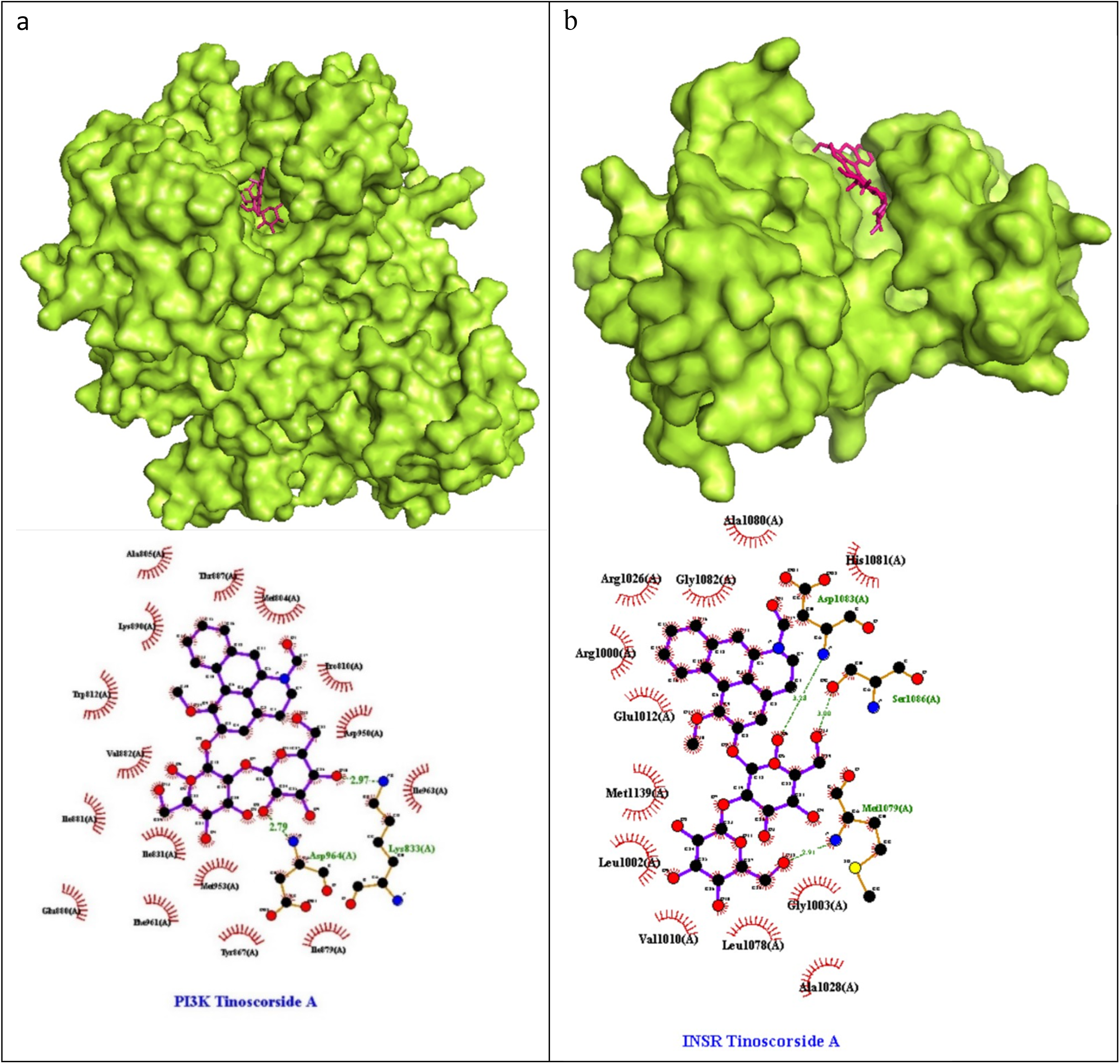
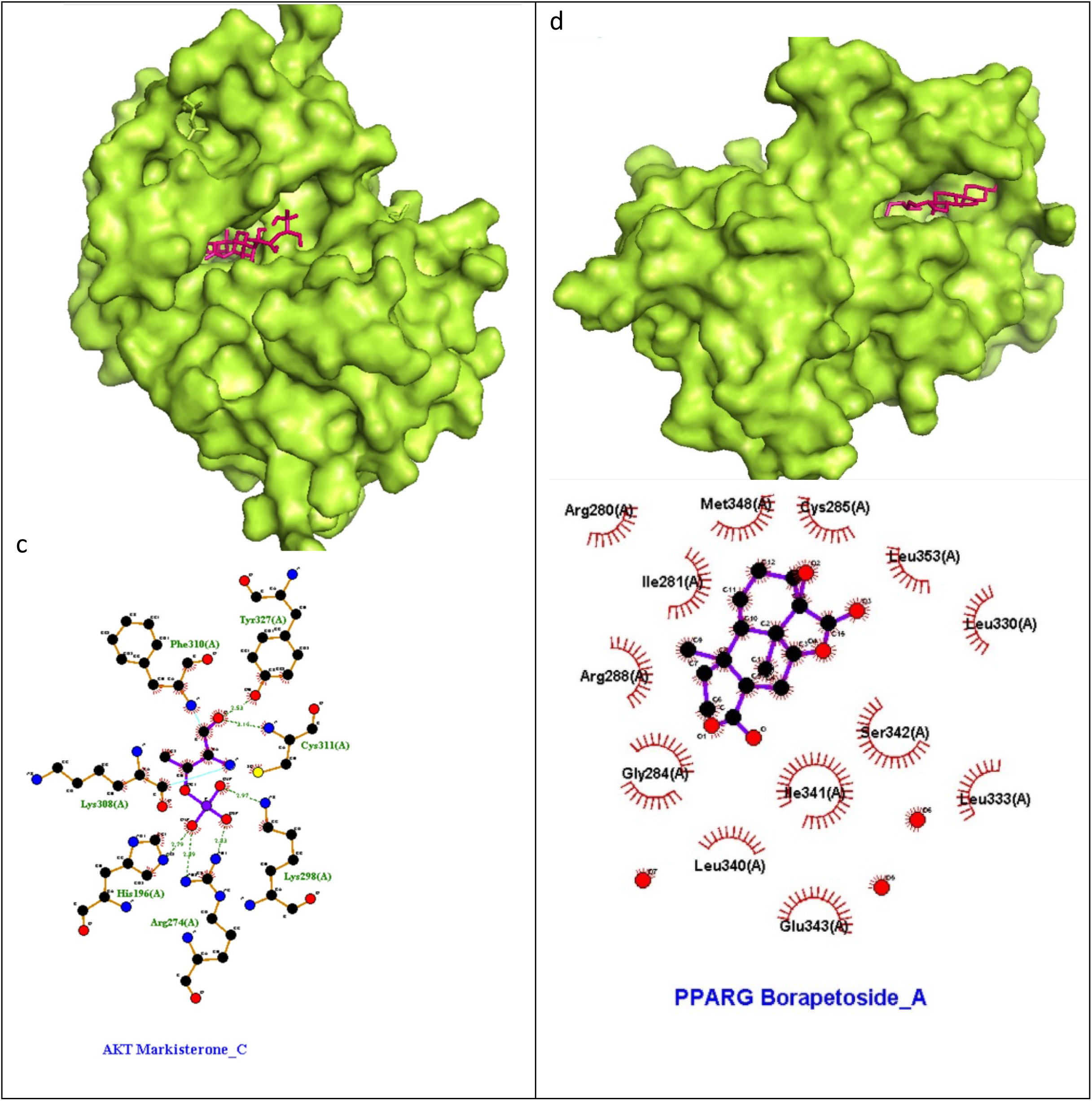
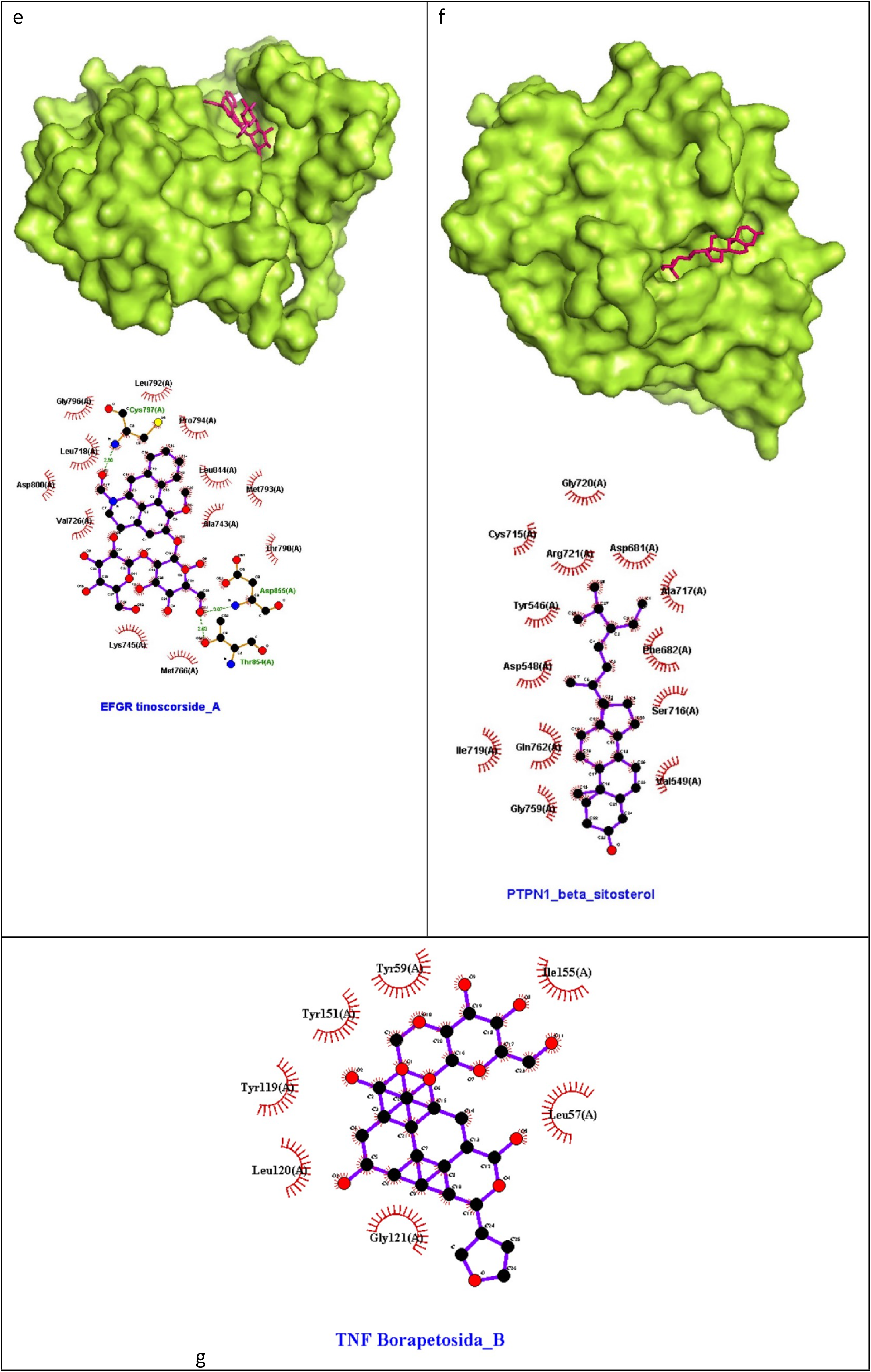
Docking sketch map of best ligand-binding position of T.crispa phytoconstituents and 7 selected taget proteins in insulin resistance.

## Notes

### Competing Interest Statement

The authors have declared no competing interest.

